# Plasticity in the functional properties of NMDA receptors improves network stability during severe energy stress

**DOI:** 10.1101/2023.01.19.524811

**Authors:** Nikolaus Bueschke, Lara Amaral-Silva, Min Hu, Alvaro Alvarez, Joseph M. Santin

## Abstract

Brain energy stress leads to neuronal hyperexcitability followed by a rapid loss of function and cell death. In contrast, the frog brainstem switches into a state of extreme metabolic resilience that allows them to maintain motor function during hypoxia as they emerge from hibernation. NMDA receptors (NMDARs) are Ca^2+^-permeable glutamate receptors that contribute to the loss of homeostasis during hypoxia. Therefore, we hypothesized that hibernation leads to plasticity that reduces the role of NMDARs within neural networks to improve function during energy stress. To test this, we assessed a circuit with a large involvement of NMDAR synapses, the brainstem respiratory network of female bullfrogs, *Lithobates catesbeianus*. Contrary to our expectations, hibernation did not alter the role of NMDARs in generating network output, nor did it affect the amplitude, kinetics, and hypoxia sensitivity of NMDAR currents. Instead, hibernation strongly reduced NMDAR Ca^2+^ permeability and enhanced desensitization during repetitive stimulation. Under severe hypoxia, the normal NMDAR profile caused network hyperexcitability within minutes, which was mitigated by blocking NMDARs. After hibernation, the modified complement of NMDARs protected against hyperexcitability, as disordered output did not occur for at least one hour in hypoxia. These findings uncover state-dependence in the plasticity of NMDARs, whereby multiple changes to receptor function improve neural performance during energy stress without interfering with its normal role during healthy activity.

**Significance Statement:** Neural circuits lose homeostasis during severe energy stress, and NMDA-glutamate receptors play a major role in this response. In contrast, frogs have the remarkable capacity to use plasticity that improves circuit function from minutes to hours during hypoxia, likely as an adaptation to survive emergence from hibernation. We found this occurs, in part, through modification of NMDA receptors that renders them less permeable to Ca^2+^ and more likely to desensitize during high activity states. These NMDA receptor modifications do not influence normal network function but protect against hyperexcitability caused by hypoxia. This work points to endogenous plasticity mechanisms that improve network function during energy stress without altering circuit function when the brain is well-oxygenated.

## Introduction

Brain function requires high rates of ATP synthesis (Bordone et al., 2019). The sudden loss of oxygen triggers hyperexcitability that leads to ion dysregulation and cell death, termed “excitotoxicity” (Buck & Pamenter, 2018). A key contributor to excitotoxicity involves the activation of NMDA-glutamate receptors (NMDARs), causing pathological Ca^2+^ influx and saturation of intracellular Ca^2+^ buffering systems (Szydlowska & Tymianski, 2010; White & Reynolds, 1995). As rates of ATP synthesis wane, active transport of Ca^2+^ out of the cell becomes increasingly difficult, which culminates in cell death through reactive oxygen species, pro-death signaling pathways, and mitochondrial damage (Wu & Tymianski, 2018). Therefore, hyperexcitability caused by overstimulation of NMDARs and subsequent Ca^2+^ influx represents a critical step in the loss of homeostasis that drives neurological issues during hypoxia.

Unlike most mammals, some species experience variable oxygen tensions in their environments and have evolved strategies to survive brain hypoxia (Larson et al., 2014). Survival strategies often involve entry into a hypometabolic state associated with arrested synaptic transmission to conserve energy (Buck & Pamenter, 2018). However, some animals must remain active during hypoxia (Czech-Damal et al., 2014; Larson et al., 2014; Larson & Park, 2009). This presents the challenge of maintaining circuit function while avoiding hyperexcitability and excitotoxicity. An extreme example of this problem is embodied by the respiratory network of American bullfrogs. This network generates rhythmic output through mechanisms that involve AMPA and NMDA-glutamate receptors (Kottick et al., 2013) and, therefore, requires ongoing aerobic metabolism (Adams et al., 2021). However, for several months each year, frogs hibernate in ice-covered ponds without breathing air, using only skin gas exchange. As a consequence, blood oxygen falls dramatically, which may be as low as 1-3 mmHg (Tattersall & Ultsch, 2008). Low oxygen in this environment does not pose an immediate threat due to reduced metabolic rates in the cold (Tattersall & Boutilier, 1997). However, during emergence at warm temperatures, this life sustaining network, must restart activity on the background of severe hypoxia. If this was not already difficult enough, an additional problem lies in the fact that frogs are not generally considered to be strongly “hypoxia-tolerant,” and low O_2_ levels present during emergence cannot power brainstem circuits (Adams et al., 2021). To overcome these challenges, hibernation induces metabolic plasticity at synapses that improve network function during severe hypoxia from a few minutes to several hours (Bueschke et al., 2021, Amaral-Silva and Santin, 2023; Hu and Santin, 2022). Therefore, hibernation in frogs provides insight into plasticity that shifts a typically “hypoxia-intolerant” circuit into a state that functions remarkably well during severe energy stress.

As in mammals, hypoxia in the frog brain induces network hyperexcitability that disrupts patterned output, followed by a swift loss of function (Adams et al., 2021; Bueschke et al., 2021a). Thus, this network must engage mechanisms that constrain excitability to maintain activity with a limited energy supply upon emergence from hibernation. Many hypoxia-tolerant vertebrates have low levels of NMDARs or suppress NMDARs in hypoxia to conserve energy and avoid excitotoxicity (Bickler et al., 2000; Bickler & Buck, 2007; Wilkie et al., 2008). Therefore, we hypothesized that shifting from “hypoxia intolerance” to “functional hypoxia tolerance” involves reduced NMDAR function within the network. To test this hypothesis, we assessed the NMDAR tone of the respiratory network, NMDAR currents using whole-cell voltage-clamp, and NMDAR subunit composition using single-cell quantitative PCR. In contrast to our hypothesis, hibernation did not influence the role of NMDARs in the network, current amplitude, kinetics, and hypoxia sensitivity. We instead found that hibernation decreased the Ca^2+^-permeability of NMDARs and enhanced desensitization, serving to reduce Ca^2+^ influx and lower depolarizing drive during high activity states. These modifications oppose the loss of homeostasis driven by the normal profile of NMDARs. Overall, we identified NMDAR plasticity that improves network activity during energy stress without influencing their contribution in well-oxygenated conditions.

## Materials and Methods

### Animal husbandry and ethical approval

The use of animals was approved by the Institutional Animal Care and Use Committee (IACUC) at the University of North Carolina at Greensboro (Protocol #19-006). Female American bullfrogs (*Lithobates catesbeianus*) were ordered from Phrog Farm (Twin Falls, ID, USA). Frogs were randomly assigned to one of two groups: control or hibernation and were placed in designated plastic tubs containing dechlorinated water bubbled with room air. Control frogs were kept at ambient room temperatures (20°C) with access to wet and dry areas and fed pellets provided by Phrog Pharm once a week. Plastic tubs containing frogs assigned for aquatic overwintering were placed into low-temperature incubators (Thermo Fisher Scientific, Waltham, MA, USA) that were gradually reduced from 20° to 4° C over the course of 7 days. Once at overwintering temperatures, screens were placed below water level to ensure frogs’ cutaneous respiration as the primary oxygen exchange, and they were kept at 4° C for 30 days before use. Frogs exhibit metabolic suppression at these temperatures, consistent with hibernation (Tattersall & Ultsch, 2008). Therefore, we refer to this group as “hibernation.” All frogs were kept under a 12:12 hour light/dark cycle.

### Dissection of Brainstem

Frogs were deeply anesthetized with isoflurane (1 mL) in a sealed 1 L container until loss of toe-pinch response. They were then rapidly decapitated, and heads were submerged for dissection in chilled artificial cerebrospinal fluid (aCSF; 104 mM NaCl, 4 mM KCl, 1.4 mM MgCl_2_, 7.5 mM D-glucose, 1 mM NaH_2_PO_4_, 40 mM NaHCO_3_, and 2.5 mM CaCl_2_; (Bueschke et al., 2021), which was bubbled with 98.5% O_2_ and 1.5% CO_2_ for oxygenation, resulting in aCSF with a pH of 7.9 ± 0.1. These CO_2_ and pH values are normal for frogs at room temperature (Howell et al., 1970). The brainstem-spinal cord was rapidly exposed, and the forebrain was crushed. Brainstem-spinal cord nerve roots were carefully trimmed, allowing its’ excision from the cranium, which was followed by the dura membrane removal. Following all dissections, the preparations were held at room temperature (22±1°C) and all experiments were performed at this temperature.

### Extracellular nerve root recordings

For experiments that assessed motor output from the intact network, we assessed extracellular motor output from the vagus nerve rootlets, which innervates the glottal dilator muscle to control airflow in and out of the lung. Dissected preparations were pinned ventral side up in 6 mL Petri dishes coated with Sylgard 184 (Dow Inc. Midland, MI, USA). Brainstem-spinal cords were continuously superfused with oxygenated aCSF using a peristaltic pump (Watson Marlow, Falmouth, CNL, UK). Population activity from the vagal nerve root (CNX) was recorded using a suction electrode attached to a fire-polished borosilicate glass pulled from a horizontal pipette puller (Sutter Instrument, Novato, CA, USA). Extracellular signals were amplified (x1000) and filtered (low-pass, 1000 Hz; high pass, 100 Hz) using an AM-Systems 1700 amplifier (Sequim, WA, USA), then digitized using Powerlab 8/35 (ADInstruments, Dunedin, New Zealand). The raw signal was integrated and rectified (100 ms τ) using LabChart data acquisition system (ADInsruments). After 4 hours of decapitation, a stable signal was recorded for ∼30 minutes, and the preparation was challenged with hypoxic aCSF (98.5% N_2_ balanced with CO_2_) or NMDAR antagonist, D-AP5 (50 µM). All preparations included in this study produced rhythmic motor output associated with lung breathing.

### Motoneuron labeling and slice preparation

For experiments that assessed NMDA receptor currents in motoneurons from brain slices, we isolated the 4^th^ root of the glossopharyngeal-vagus complex and attached a fire-polished borosilicate glass pipette to it. Tetramethylrhodamine dextran dye was then added to the tip of the pipette (Invitrogen, Waltham, MA, USA) in contact to the nerve root for at least 2 hours to diffuse to the soma. We then sliced the brainstem-spinal cord preparations at 300 μM using a vibratome (Technical Products International series 1000, St Louis, MO, USA). During dye loading and slicing the tissue was maintained in regular aCSF (described above).

### NMDAR Currents

Slices were superfused with an extracellular solution (76.5 mM NaCl, 2.2 mM KCl, 7.5 mM D-Glucose, 10 mM HEPES, 300 μM CdCl_2_, 20 mM TEA-Cl, 250 nM TTX, 10 μM DNQX) containing two different Ca^2+^ concentrations. High Ca^2+^ had 10 mM of CaCl_2,_ and low Ca^2+^ 1 mM of CaCl_2_. Sucrose was added to maintain consistent osmolarity (330 mOsm). Thus, 60mM was added to the high Ca^2+^ solution, and 89mM was added to the low Ca^2+^ solution. MgCl_2_ was excluded to prevent Mg^2+^ block of NMDARs. Extracellular solutions were oxygenated by bubbling 98.5% O_2_ and 1.5% CO_2_.

Labeled vagal neurons were approached by glass pipettes (2–4 MΩ resistance) using a micromanipulator (MP-285/ MPC-200, Sutter Instruments, Novato, CA, USA) attached to a head stage (CV203BU, Molecular Devices, San Jose, CA, USA). Positive pressure was applied to the pipette while approaching the cell and quickly removed, gentle negative pressure was used to form a >1GΩ, and the whole-cell access was obtained by breaking the seal with rapid negative pressure. We did not observe any obvious differences in cell viability and ability to obtain patch clamp recordings in high and low Ca^2+^ solutions. The solution filling the patch pipette (76.5 mM K-gluconate, 10 mM D-glucose, 10 mM HEPES, 1 mM Na_2_-ATP, 0.1 mM Na_2_-GTP, 2 mM MgCl_2_, 30 mM TEA-Cl) was designed with Na^+^ and K^+^ concentrations equal and opposite to the extracellular solution to generate a reversal (E_rev_) potential of ∼0 mV without Ca^2+^(assuming no difference in ion selectivity between Na^+^ and K^+^) as seen previously (Jatzke et al., 2002).

To evoke NMDAR currents, we focally applied NMDA and glycine to the cell body. For this, a borosilicate pipette pulled with a tip diameter of ∼5 μm was filled with NMDA (1 mM) and glycine (10 μM) (both from Hellobio, Princeton, NJ, USA) dissolved in the extracellular solution. The pipette wasdriven by a Picospritzer II, and the focal solution was applied during 10-20 ms onto the soma of labeled vagal motoneurons (General Valve Corporation, Fairfield, NJ, USA) to activate NMDARs. Glycine is a co-agonist of the NMDAR but did not elicit a response when applied alone in control or hibernation neurons.

NMDAR E_rev_ was recorded by a voltage clamp step protocol with a Δ+5 mV step between -20 mV and 40 mV. In 3 recordings, E_rev_ was more hyperpolarized, and those cells were stepped from -40 mV to 20 mV in 5 mV increments (control, n=16; hibernation, n=16). NMDAR desensitization was assessed by puffing at a rate of 0.5 Hz for a total of ten pulses at -20 mV (control, n=12; hibernation, n=14). Hypoxia was applied to brain slices by bubbling extracellular solution with 98.5% of N_2_ and 1.5% of CO_2_ for ∼5 minutes before recording. All desensitization protocols and single activation amplitude and kinetics experiments were performed on neurons held at -20 mV. All data were acquired in pClamp 11 software using Axopatch 200B amplifier and Axon Digidata 1550B digitizer (Molecular Devices, San Jose, CA, USA).

### NMDAR subunits electrophysiology

Vagus motoneurons were recorded using patch-clamp as described above. For this protocol, the pipette solution contained 110 mM K-gluconate, 2 mM MgCl_2_, 10 mM HEPES, 1 mM Na_2_-ATP, 0.1 mM Na_2_-GTP, and 2.5 mM EGTA. Once gaining whole cell access the solution bathing the slice was changed from regular aCSF to 104 mM NaCl, 4 mM KCl, 7.5 mM D-glucose, 1 mM NaH_2_PO_4_, 40 mM NaHCO_3_, 2.5 mM CaCl_2_, 250 nM TTX, 10 µM DNQX, 3 µM Glycine, and 2 µM Strychnine. NMDA currents were elicited once every minute by focal application of NMDA (1 mM) and glycine (3 μM) diluted in the aCSF in 500 ms pulses. The cell was continuously monitored in voltage clamp, and after observing a stable current in 3 consecutive NMDA puffs (∼ 10-15 min of recording), we applied a specific NMDA subunit inhibitor in aCSF.

The subunit GluN2A was inhibited using 1 µM TCN 201 (Tocris Bioscience, Bristol, UK) (Bettini et al., 2010; Edman et al., 2012); GluN2B was inhibited by 2 µM RO 25-6981 (Tocris Bioscience, Bristol, UK) (Abrahamsson et al., 2017; France et al., 2017); GluN2C/GluN2D was inhibited using 10 µM QZN 46 (Tocris Bioscience, Bristol, UK) (Hansen and Traynelis, 2011); and GluN3 was blocked by 30 µM TK30 (4-(2,4-dichlorobenzoyl)-1H-pyrrole-2-carboxylic acid; Santa Cruz Biotechnology, Dallas, TX, USA) (Kvist et al., 2013; Christian et al., 2021). A time control experiment was performed recording NMDAR currents while maintaining the slice in the aCSF described above with no inhibitor added.

### Electrophysiology Data analysis

#### Extracellular motor output

Burst amplitude and frequency were determined by averaging amplitude of fictive breaths selected in a 5-minute window prior to D-AP5 application. Amplitude and frequency in the presence of NMDAR antagonist were analyzed for 5 minutes after burst amplitude stabilized. Peak amplitude was determined using the Peak Analysis extension in LabChart (ADInstruments, Dunedin, New Zealand). To characterize chaotic bursting behavior during hypoxia, we analyzed the peak of the largest “non-respiratory” motor burst that clearly disrupted patterned network output associated with respiratory activity and normalized it to the background nerve signal value at the baseline to allow comparisons across groups.

#### NMDAR Ca^2+^ permeability

Relative Ca^2+^ permeability was determined by the degree of shift in the NMDAR reversal potential (E_rev_) based on the mean data of the population in different Ca^2+^ concentrations in extracellular recording solution. The underlying premise is that receptors that are permeable to Ca^2+^ have a depolarizing shift in E_rev_, while Ca^2+^-impermeable channels do not (Jatzke et al., 2002). We evoked NMDAR currents using low (1 mM) and high (10 mM) Ca^2+^ and measured the peak current evoked by NMDA-glycine at each voltage step. We then plotted the current-voltage relationship and interpolated E_rev_ through the x-intercept of the line of best fit from 3-5 plot points near the intercept where I=0 pA. The currents elicited by NMDA-glycine were small near E_rev_ and were unlikely to be affected by voltage errors due to the series resistance (R_s_). However, the holding current at depolarized voltages was often substantial, even in the presence of TEA to block outward currents. Therefore, we corrected the holding voltage for R_s_ errors. For this, we measured R_s_ at each voltage step based on the peak of the transient current and corrected the holding voltage by the voltage error caused by R_s_. Currents were measured using the Peak Analysis extension in LabChart. Averaging/decimation at 0.05-0.1 ms was applied to the traces prior to peak analysis to filter out high-frequency spontaneous synaptic activity. This level of filtering was chosen as it did not alter the amplitude of the NMDAR current.

#### Single NMDAR currents and desensitization

Single NMDAR currents were measured using similar methods as the NMDAR Ca^2+^ permeability, where peak currents induced by NMDA and glycine were assessed at -20 mV. Decay time constant was determined between 90% and 10% of the peak height and calculated using the Peak Analysis tool in LabChart. Desensitization was assessed by measuring the peak current evoked at each step. The baseline for each current in the series was taken as the recovered current after proceeding puff.

NMDAR subunit inhibition: The current amplitude was analyzed using peak analysis in LabChart. The average of the last 3 currents before inhibitor application was compared to currents recorded after 10 minutes of exposure to the drug. A set of time control experiments compared the 3 first stable currents (parameter used to decide to apply the inhibitor) to 10 min after that.

### Single-cell Real-time quantitative PCR

We used single-cell quantitative PCR (qPCR) to determine mRNA expression for NMDARs subunits using the same cell harvesting, RNA extraction, cDNA synthesis, and preamplification procedures detailed in Pellizzari et al., 2023.

PCR primers for open reading frames of the genes that code for NMDA glutamate receptor subunits, *Grin1, Grin2a, Grin2b, Grin2c, Grin2d, Grin3a, and Grin3b,* were designed from sequences found in the coding DNA sequence for *Lithobates catesbeianus* (Table 1 and Table 2). For this, we used annotated amino acid sequences for GluN subunits from *Rana temporaria* as a query in the *Lithobates catesbeianus* amino acid database. This search revealed peptide sequences with high amino acid sequence conservation. We then performed a reciprocal BLAST against the entire nonredundant protein database using hits from *Lithobates catesbeianus* to verify the identity of the target. Accession numbers were then used to identify the open reading frame in the CDS to design PCR primers. Only *Grin1, Grin2a, and Grin3a* were identified in the bullfrog CDS, likely due to low coverage and/or poor assembly of the *L. catesbeiansus* genome (Hammond et al., 2017). For *Grin2b, Grin2c, Grin2d, and Grin3b*, we found the regions of the coding sequence in *Rana temporaria* (a species closely related to *L. catesbeianus*) and *N. parkeri* (more distantly related frog species) with high similarity to design PCR primers. Our rationale was that close sequence identity at the nucleotide level between these two species (∼98%) would allow us to design primers for use in *Lithobates catesbeianus*.

**Table 1.**
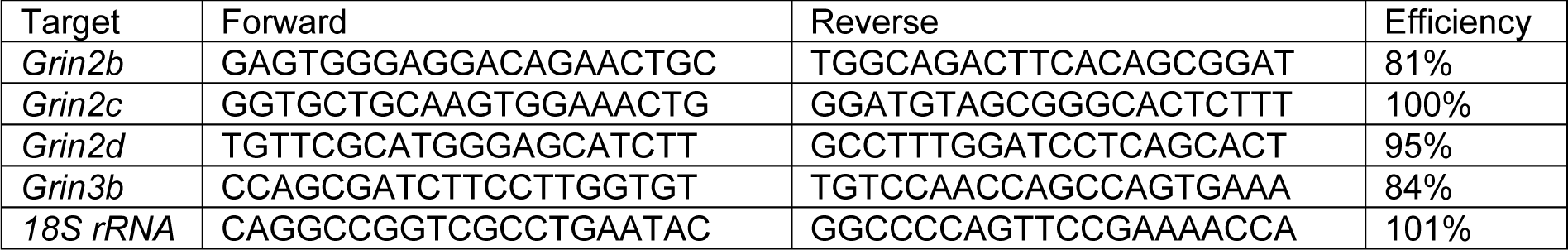
Primer Sequences for SYBR Green qPCR Assays.

**Table 2.**
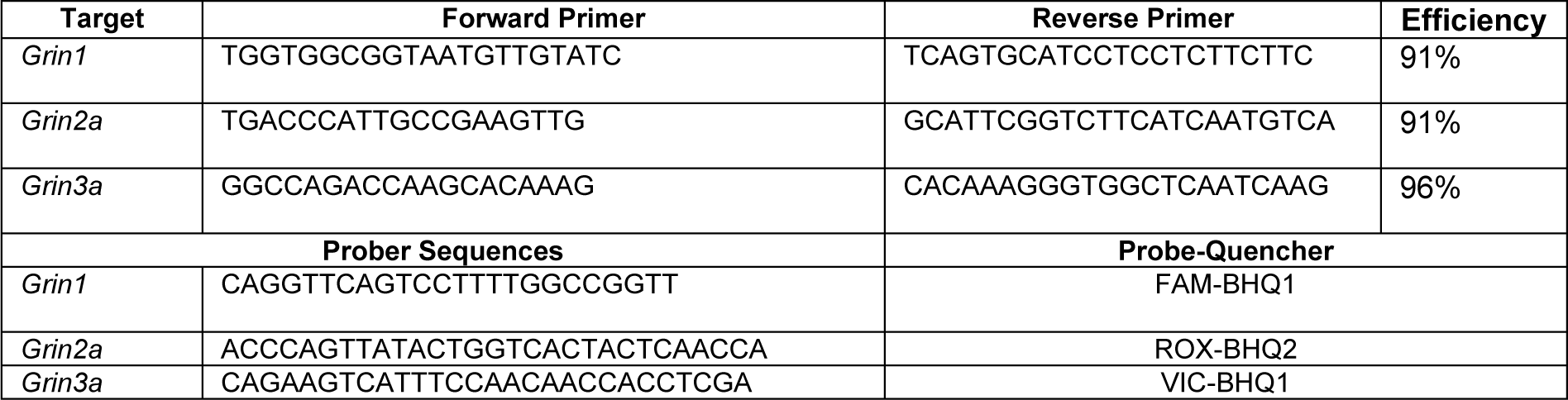
Primer Sequences for Probe-based qPCR Assays.

Primer sets were validated with a series of four four-fold dilutions of brainstem cDNA. All primer sets used here produced efficiencies greater than 80% (Table 1 and Table 2) and a single peak in the melt curve in a SYBR Green assay, suggesting the amplification of a single PCR product. As we observed in Pellizzari et al., 2023 some primer sets that produced one peak in the melt curve using bulk brain cDNA as the input material showed multiple peaks in the melt curve after single-cell preamplification. When this occurred, these primer sets were redesigned.

All neurons in this experiment were assessed for the expression of each of the 7 *Grin* subtypes. *Grin2b, Grin2c, Grin2d, and Grin3b* were run using SYBR Green assays, and *Grin1, Grin2a, and* Grin3a were run in one multiplex assay. For SYBR Green assays, quantitative PCR was run in 10 µL reaction volumes containing 2.5 µM forward and reverse primers and followed the instructions of the 2X SYBR Green Mastermix (Applied Biosystems, ThermoFisher Scientific, Waltham, MA). Assays were run on 96-well plates on an Applied Biosystems QuantStudio 3 (Applied Biosystems, ThermoFisher Scientific, Waltham, MA) using the following cycling conditions according to the SYBR Green instructions: 50°C-2m, 95 °C-10 m, 95 °C-15 s, 60 °C-1 m. Following 40 cycles of PCR (95°C-15 s, 60°C-1 m), melt curves for all PCR products were acquired by increasing the temperature in increments of 0.3°C for 5 seconds from 60°C to 95 °C. For multiplex assays, we ran triplexed probed-based assays. For this, we used the same primer concentration as described for SYBR Green assays, 312.5 nM reporter probes, and followed the instructions of the 5X PerfeCTa qPCR Toughmix mastermix (Quanta Bio). 18s ribosomal RNA was run to ensure the quality of the sample and for normalization of copy number to account for the possibility of different amounts on cDNA input and efficiency in the cDNA synthesis reaction.

Absolute quantitation of transcript abundance was estimated through copy number standard curves as previous described (Santin & Schulz, 2019). We normalized absolute copy number by a normalization factor using 18s Cq values to account for the possibility of different amounts of harvested cytoplasm and efficiencies of the cDNA synthesis reaction across samples (Garcia et al., 2018). We ran qPCR for all *Grin* subunits on 19 control neurons and 20 hibernation neurons. In some neurons, *Grin* subunits appeared to be absent, or to be very lowly expressed. However, it is also possible a lack of detection represents a false negative due to stochasticity in the cDNA synthesis reaction due low RNA input quantities associated with the single cell. Thus, we included *Grin* genes in analysis that had C_q_ values <29.5 after preamplification. The only exception to this was for *Grin2D.* The abundance was consistently low for most samples; therefore, we included all data points in the analysis for *Grin2D*.

### Statistical analysis

Statistical analysis was performed using GraphPad Prism 9 (GraphPad Software, San Diego, CA, USA). Comparisons between independent 2 groups were carried out with a two-tailed unpaired t-test. With 3 or more groups, we used a one-way ANOVA. In experiments with a “before-after” design, we used a two-tailed paired t-test. In experiments with two main effects, we used a two-way ANOVA. One-way and two-way ANOVA were followed up with Holm-Sidak multiple comparison test when appropriate. Individual data points were presented in addition to mean±SD or with box-and-whisker plots, on the latter, boxes represent the interquartile range, and whiskers indicate maximum and minimum values in the data set. Significant was accepted when p<0.05.

## Results

Function of the brainstem respiratory network in severe hypoxia increases by hours after animals emerge from hibernation (Bueschke et al., 2021; Amaral-Silva & Santin, 2023). To determine if reductions in NMDAR function play a role in this response, we assessed the NMDAR tone in *in vitro* brainstem-spinal cord preparations. This preparation produces rhythmic output associated with breathing that can be recorded from the cranial nerve X rootlet (Fig. 1A). NMDARs are involved in transmitting synaptic input from the respiratory rhythm generator to the motor pools (Kottick et al., 2013). If the contribution of NMDAR was reduced following hibernation, we expected to observe a lower sensitivity to the block of NMDARs using D-AP5 relative to controls. D-APV reduced the amplitude of the motor output by ∼50%, corroborating previous results demonstrating that NMDAR transmission plays a large role in recruiting motoneurons to the population burst (Kottick et al., 2013). However, sensitivity of the motor amplitude to D-APV did not change after hibernation (Fig. 1B-C, t_(13)_ = 0.9325, p = 0.37, unpaired t-test). In addition, sensitivity of respiratory burst frequency to D-APV, which provides an assessment of rhythm generator function, did not change after hibernation (Fig. 1D, t_(13)_ = 1.699, p = 0.11, unpaired t-test). These results show that the overall contribution of NMDARs to network function is unchanged following hibernation.

**Figure 1.**
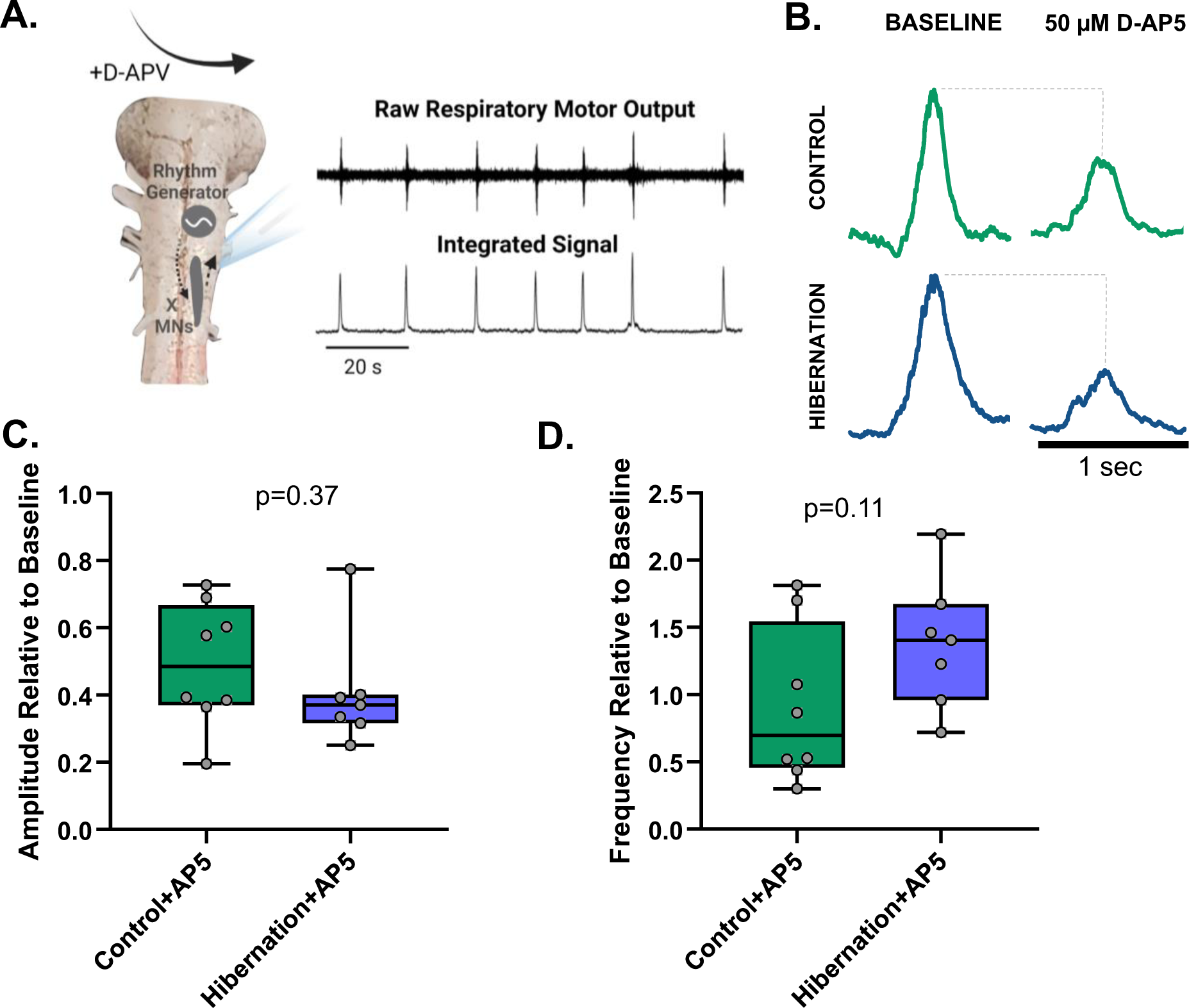
Hibernation does not alter the NMDAR tone of respiratory motor output. (A) Schematic of the in vitro brainstem preparation. Raw motor output associated with breathing can be recorded from the cut cranial nerve X rootlet (CNX). Raw signals are then rectified and integrated for analysis. (B) Representative respiratory motor bursts in baseline conditions and after block of NMDA-glutamate receptors with 50 µM D-AP5. Control animals are shown on the top and hibernation is shown on the bottom. (C) Box and whisker plot comparing burst amplitude in D-AP5 relative to control between controls (left, green, n = 8) and hibernation (right, blue, n = 7) brainstems, showing no significant change in D-AP5 sensitivity (unpaired t-test). (D) Box and whisker plots comparing burst frequency changes in response to D-AP5. There was no significant difference between the change in burst frequency induced by D-AP5 across groups. Dots represent each data point of individual experiments.

Although the contribution of NMDAR to network function did not change, we hypothesized that functional properties of the NMDAR may play a role in improving robustness in hypoxia. For example, plasticity in current amplitude or kinetics (*i.e*., smaller currents with fast deactivation times) may lower ionic fluxes through NMDARs, and, therefore, play a role in energy conservation. Moreover, some hypoxia-tolerant species contain NMDARs that are inhibited by hypoxia, offsetting excitotoxicity during energetic stress (Bickler et al., 2000). Therefore, we measured NMDARs currents in identified vagal motoneurons from brain slices using whole-cell voltage clamp in response to focal application of the NMDAR co-agonists, 1 mM NMDA and 10 µM glycine (Fig. 2A). Current amplitude and deactivation time constants at -20 mV did not change in response to hibernation (Fig 2B-D, amplitude: t_(24)_ = 1.502, p = 0.1461, unpaired t-test; deactivation time constant: t_(24)_ = 0.4516, p = 0.6556, unpaired t-test). In addition, superfusion of aCSF bubbled with a hypoxic gas mixture (0% O_2_) for 5 minutes did not consistently alter the amplitude or deactivation time in controls (Fig 2E, amplitude: t_(4)_ = 1.455, p = 0.2193, paired t-test; deactivation time constant: t(4) = 1.980, p = 0.1189, paired t-test) or following hibernation (Fig 2F, amplitude: t_(3)_ = 0.2362, p=0.8285, paired t-test; deactivation time constant: t_(3)_ = 0.1515, p=0.8892, paired t-test). Therefore, hibernation does not alter whole-cell NMDAR currents, as well as their deactivation kinetics and sensitivity to acute hypoxia.

**Figure 2.**
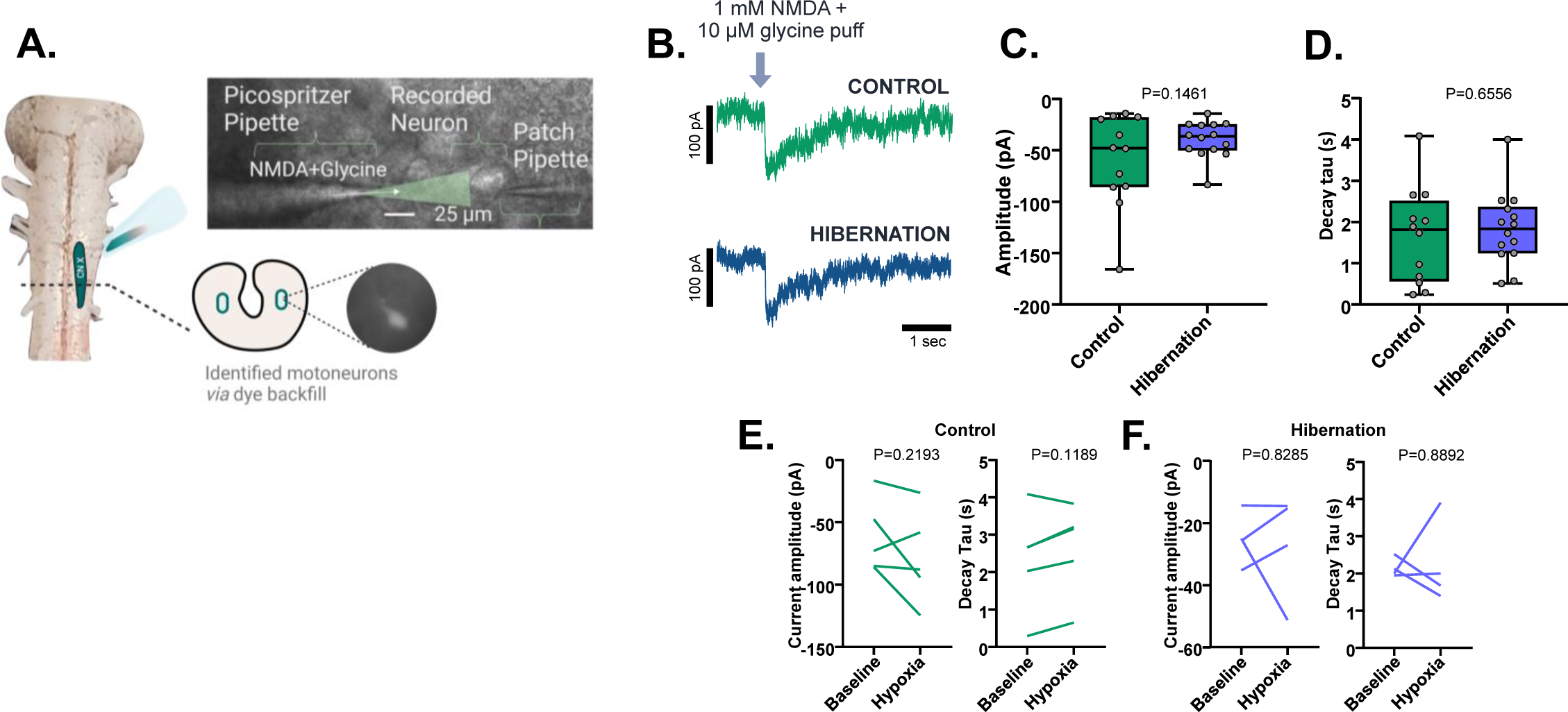
Hibernation does not alter the whole-cell NMDAR current amplitude, decay kinetics, and hypoxia sensitivity in identified respiratory motoneurons. (A) Schematic illustrating the backfill labeling process to identify motoneurons and the setup to focally apply NMDAR agonists (1 mM NMDA and 10 µM glycine) to the cell body. (B) Traces showing typical responses to focal application of NMDA and glycine. (C-D) NMDAR current amplitude and decay time constant is unaltered between control (n = 11 neurons from N = 5 animals) and hibernation (n = 13 neurons from N = 4 animals) neurons following unpaired t tests. Boxes show interquartile range and whiskers represent min and max values with dots showing individual data points. (E-F) Individual NMDAR amplitude and decay values before and after ∼5 minutes of hypoxia (0% O_2_). Paired t test results suggest hypoxia does not change either NMDAR variable in control (n = 5 neurons from N = 4 animals) or hibernation (n=4 neurons from N = 4 animals) neurons. Line represents the “before” and “after” response for individual neurons in response to hypoxia.

Ca^2+^ influx through NMDARs plays a role in network dysfunction during hypoxia (Szydlowska and Tymianski, 2010). Thus, we assessed the Ca^2+^ permeability of NMDARs before and after hibernation, as Ca^2+^ selectivity may increase or decrease without obvious changes in the whole-cell current amplitude and kinetics (Murphy et al., 2014; Skeberdis et al., 2006). For this, we measured the reversal potential of the NMDAR current (E_rev_) in low Ca^2+^ (1 mM) and high Ca^2+^ (10 mM). Changes in E_rev_ during exposure to high Ca^2+^ concentrations provide a way to assess the relative Ca^2+^ permeability of ligand-gated ion channels (Jatzke et al., 2002). Consistent with the canonical role of NMDAR as a Ca^2+^ permeable channel, raising extracellular Ca^2+^ depolarized E_rev_ of in control neurons (Fig. 3A top). Divalent cations can adhere to the outer edge of cell membranes and alter the surface charge, making the apparent voltage at the membrane more depolarized with increasing concentrations of divalent cations (Hille, 2001). This is important to consider, given that we found a depolarizing shift in E_rev_ in response to high Ca^2+^. However, charge screening is unlikely to affect our interpretation since E_rev_ shows a <2 mV difference between 1 mM and 10 mM Ca^2+^ on Ca^2+^-impermeable kainite receptors (Jatzke et al., 2002). Thus, most of the ∼13 mV depolarization we observe between low and high Ca^2+^ in control neurons likely reflects its true Ca^2+^ permeability and not changes in the surface charge. After hibernation, the change in E_rev_ during high Ca^2+^ appeared to be far smaller (Fig. 3A bottom). Indeed, there was a significant interaction between Ca^2+^ concentration and group on E_rev_ (Fig. 3C, F_(1,60)_ = 7.735, p = 0.0072, two-way ANOVA), with a strong increase in E_rev_ by high Ca^2+^ in the control group and no significant change in E_rev_ in high Ca^2+^ after hibernation. Therefore, hibernation leads to a reduction in the Ca^2+^ permeability of the NMDAR, with minimal influence on the whole-cell current amplitude and deactivation kinetics.

**Figure 3.**
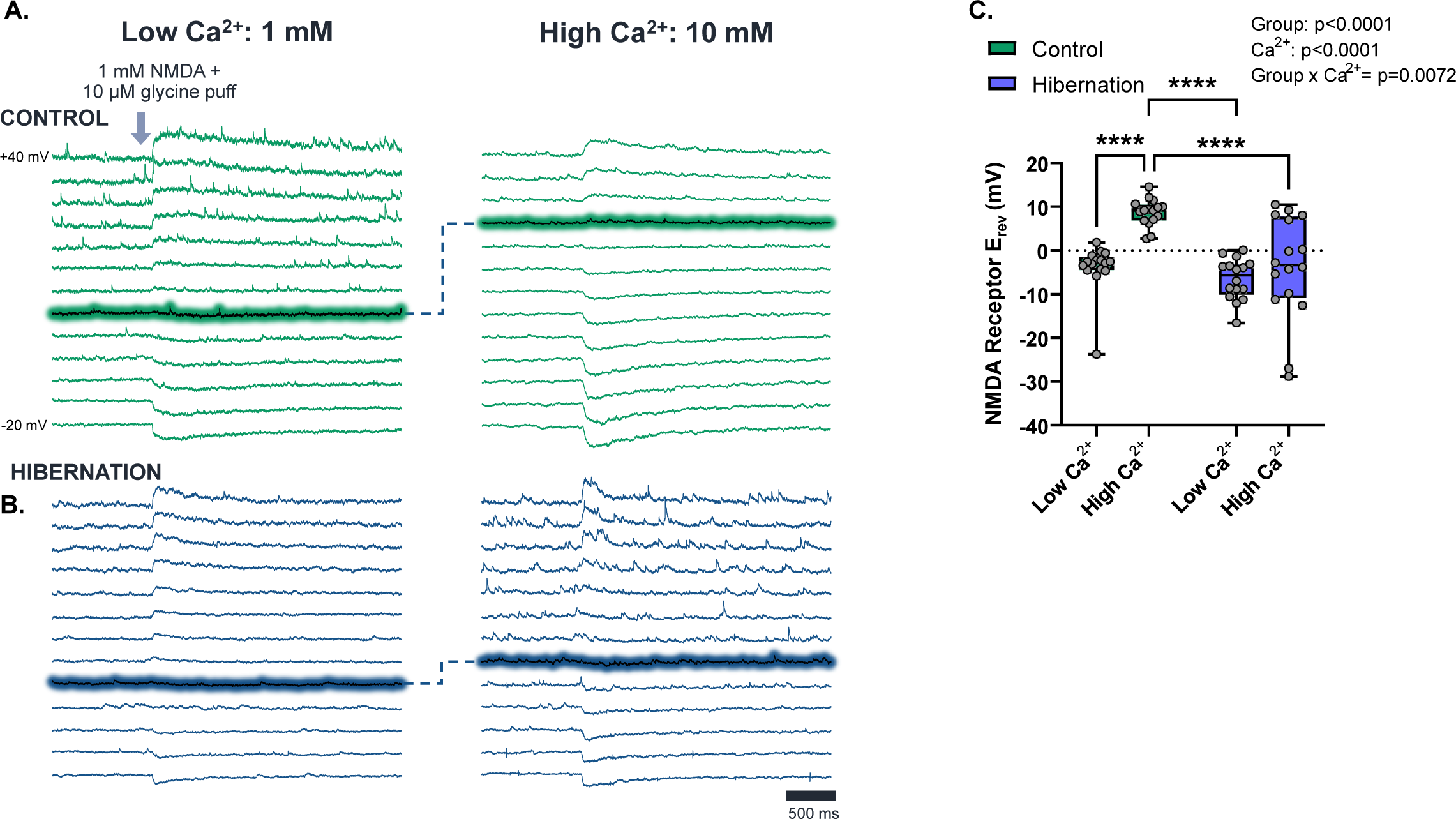
Hibernation decreases Ca^2+^ permeability of NMDARs. (A-B) Raw whole-cell voltage clamp traces held at a range of voltages (-20 to +40 mV, Δ5 mV) responding to NMDAR activation *via* focal application of NMDAR agonists. Control (A) is shown on the top in green, and hibernation (B) is shown in blue. Reversal potentials (E_rev_) in these example traces are highlighted to visualize the typical reversal potentials in 1 mM and 10 mM Ca^2+^ from control and hibernation motoneurons. (C) Results of NMDAR E_rev_ (n = 16 neurons, all groups) show significance in 2-way ANOVA interaction (p = 0.0072), indicating that hibernation affects the response to Ca^2+^. Pairwise comparisons reveal a significant depolarization of E_rev_ by 10 mM Ca^2+^ in control neurons but not hibernation neurons. We used 4 animals for hibernation in low Ca^2+^ and 5 animals for all other groups. Dots represent each data point of individual cells in each group, with boxes representing interquartile range and whiskers displaying min and max E_rev_ values. E_rev_ values shown in this plot are corrected for calculated series resistance errors based on the holding current. ****signifies p<0.0001

We next investigated the dynamic properties of NMDAR currents. Desensitization describes the degree to which a receptor loses responsiveness to the agonist in response to continued exposure, which may reflect a mechanism to dynamically reduce NMDAR currents when the network is in a high activity state. For this, we simulated NMDAR receptor activation that likely occurs during large phasic motor activation that disrupts normal rhythmic output in severe hypoxia (Fig. 4A) (Adams et al., 2021). Thus, we applied NMDAR agonists every 2 seconds for a total of 10 pulses per neuron to assess desensitization in a physiologically-relevant way (Fig 4B). We observed a moderate degree of desensitization in controls, whereby the 10^th^ pulse in the series produced a current that was ∼60% of the baseline amplitude. NMDARs from hibernators also desensitized, but the final puff elicited a current that was significantly smaller than controls, at ∼20% of the initial value (Fig. 4C, t_(24)_ = 2.946, p=0.0071, unpaired t-test). In addition, a two-way ANOVA for relative current amplitude across all puffs shows a main effect of group, indicating that currents from hibernators were more desensitized across the entire experimental protocol (Fig. 4D, F_(1, 24)_ = 7.084, p=0.0137, two-way ANOVA). Therefore, NMDAR currents not only become less permeable to Ca^2+^ after hibernation, but they also pass less current during repetitive activation that otherwise causes network output to lose homeostasis.

**Figure 4.**
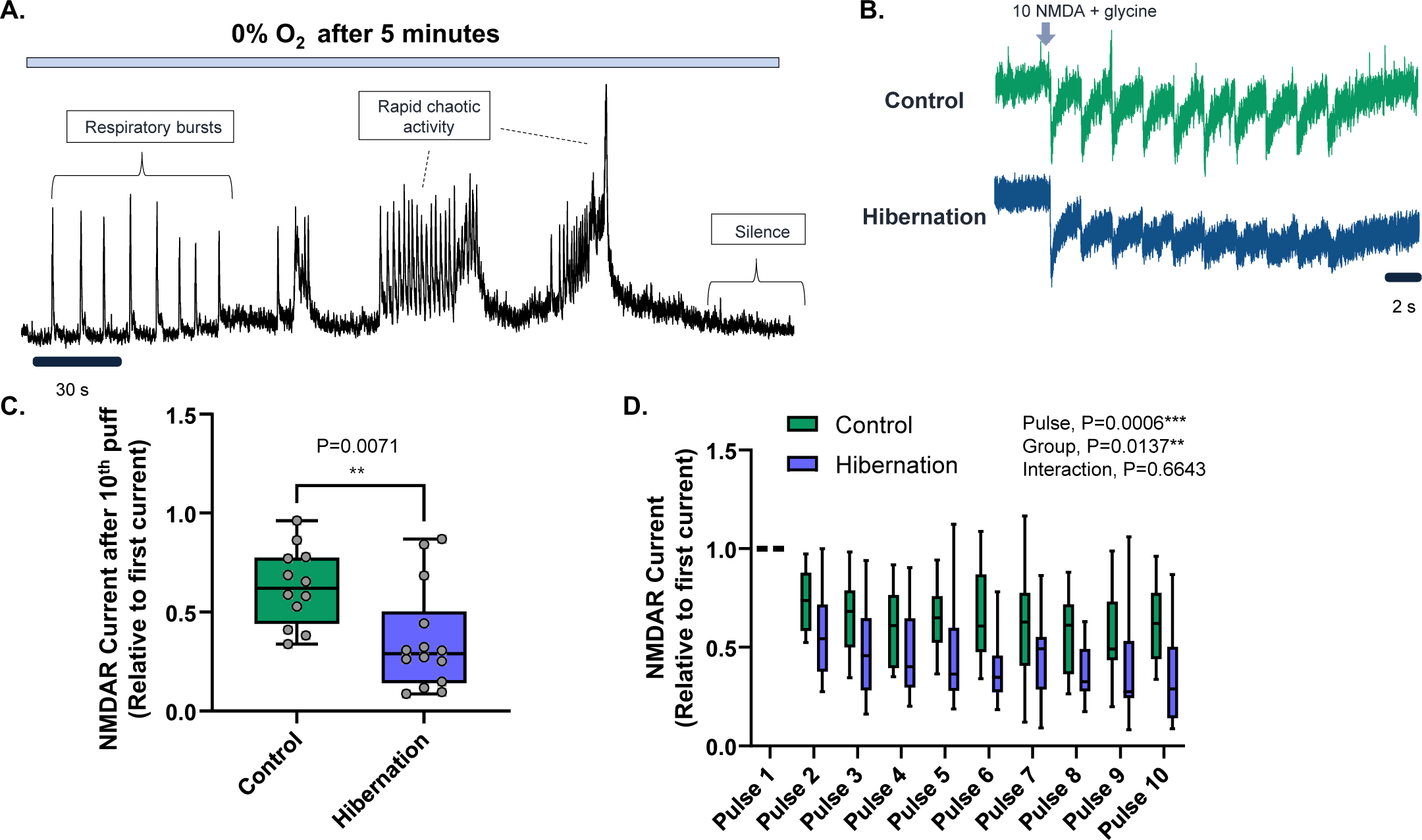
Hibernation enhances NMDAR desensitization in response to repetitive activation. (A) Example recording of population motor output from a control brainstem preparation ∼5 minutes into hypoxia. This illustrates the transition from normal patterned output to chaotic, higher frequency output with large amplitude “non-respiratory” bursting activity that occurs during hypoxia. (B) Representative traces of vagal motoneuron response to pulses of NMDAR/glycine onto the soma at 0.5 Hz. Controls under a moderate degree of desensitization (green, top), while hibernation neurons (bottom, blue) tended to have smaller current amplitudes by the end of the experiment. (C) Final (10^th^) pulse current amplitude relative to initial pulse. Hibernation motoneurons (n = 14 neuron from N = 4 animals) had a significantly lower final amplitude than controls (n = 12 neurons, N = 5 animals), suggesting that NMDARs after hibernation are more susceptible to desensitization. (D) Plot showing all data from the experimental series. Two-way ANOVA reveals a main effect of group and puff number. A significant group effect indicates that NMDAR currents throughout the experimental series were overall smaller after hibernation, consistent with enhanced desensitization. Dots represent each data point from individual cells, boxes represent the interquartile range, and lines display mean with min and max values.

NMDARs are heterotetramers composed of an obligatory GluN1 subunit and a combination of GluN2A-D and/or GluN3A-B protein subunits (Paoletti et al., 2013). To gain insight into the NMDAR subunits that correspond these physiological changes, we used inhibitors for the subunits GluN2A, GluN2B, GluN2C/GluN2D, and GluN3 to uncover the physiological participation of NMDA subunits in neurons from controls and hibernators. In controls, we observed a significant inhibition of the total NMDAR current when the GluN2B and GluN2C/GluN2D subunits were inhibited compared to time control, shown by a current decrease of 44% for GluN2B (t_17_=4.915, p=0.0001 t-test) and 29% for GluN2C/D (t_15_=2.916, p=0.0106 t-test). It is important to appreciate that time controls underwent a slight increase in current amplitude during the experimental time course (Fig 5A); thus, the decreases induced by each drug represent an underestimation of their true contribution to the total current. After hibernation GluN2B maintained its participation in the NMDA current, showing a decrease relative to time control when inhibited (t_22_=3.103, p=0.0052 t-test), which did not differ from control neurons (t_20_=1.427, p=0.1689 t-test; Fig. 5C). However, GluN2C/GluN2D no longer contributed to the NMDA current after hibernation (t_19_=0.2833, p=0.7800 t-test compared to time control), showing a difference from the control group (t_15_=2.612, p=0.0196 t-test, Fig. 5D). The subunits GluN2A (Fig. 5B) and GluN3 (Fig. 5E) did not seem to have a consistent functional contribution to the NMDA current in control or overwintered frogs. Therefore, the GluN2B subunit appears to be the main GluN2 subunit participating in the NMDA current, while GluN2C/D plays a smaller role and is lost after hibernation.

**Figure 5.**
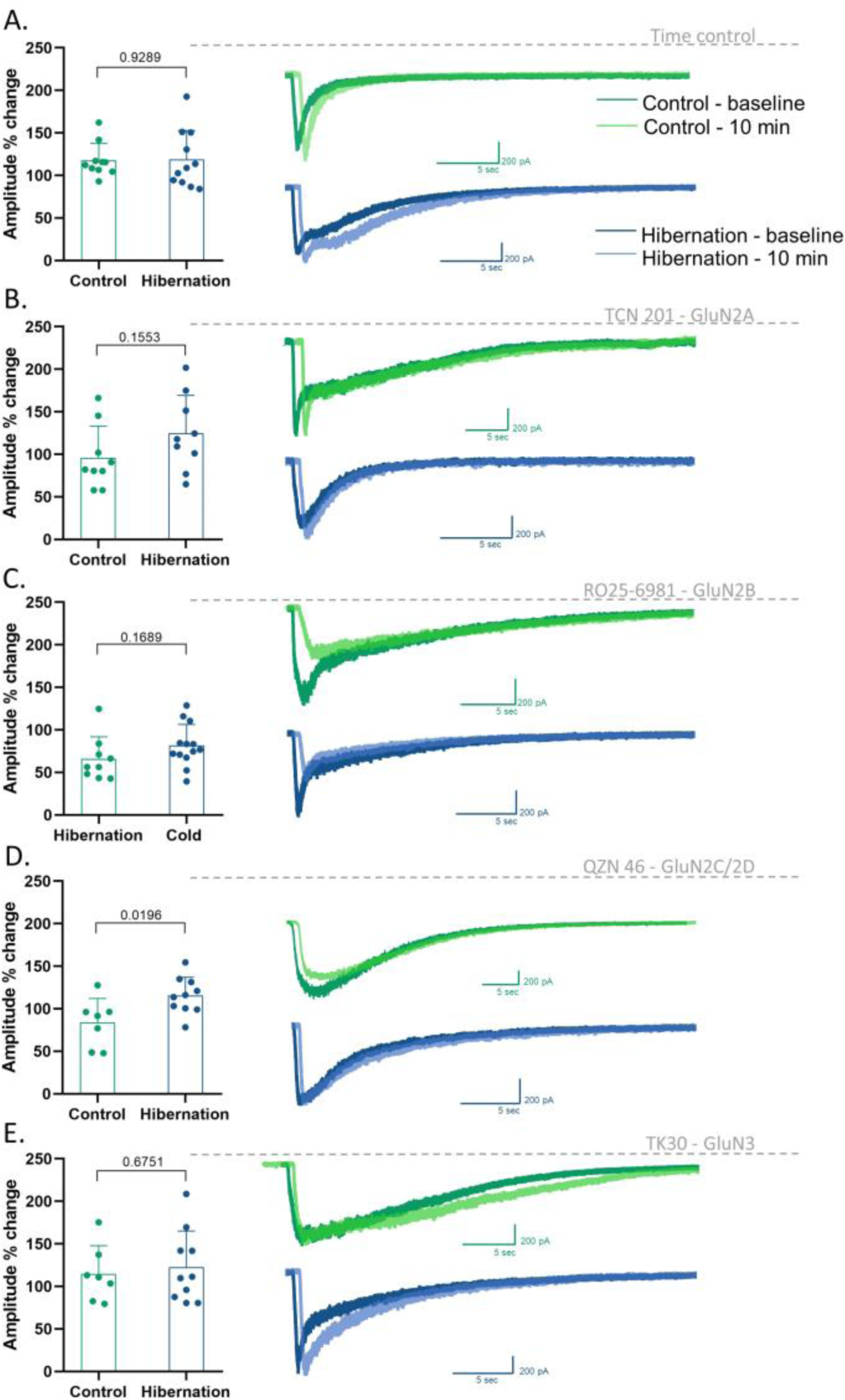
Physiological contributions from NMDAR subunits and changes after hibernation. Change in NMDAR current amplitude after 10 minutes of recording (A, Time control), inhibition of GluN2A (B, 1 µM TCN 201), GluN2B (C, 2 µM RO 25-6981), GluN2C/GluN2D (D, 10 µM QZN 46), and GluN3 (E, 30 µM TK30). Left - change in current amplitude (expressed as % of baseline) in control neurons compared to after hibernation. Right-Example traces of a baseline current overlapped with a current after 10 min of recording (A) or after 10 min of the inhibitor exposure (B-E) in neurons from frogs in control and hibernation conditions. The “after drug” or time controls traces are slightly offset from their respective baseline recordings to enhance visibility. Number of cells (n) and frogs (N) used in these experiments: Time control in control n= 10, N= 4, in hibernation n=11, N=7; GluN2A inhibition in control n= 9, N= 3, in hibernation n=9, N=6; GluN2B inhibition in control n= 9, N= 3, in hibernation n=13, N=6; GluN2C/GluN2D inhibition in control n= 7, N= 3, in hibernation n=10, N=6; GluN3 inhibition in control n= 7, N= 2, in hibernation n=10, N=9.

To understand the potential for transcriptional control of NMDAR subunits, we followed up physiology experiments with single-cell quantitative PCR (qPCR) to measure the mRNA expression of all 7 NMDAR subunits that encode the NMDAR (the *Grin* gene family). These data are summarized in Fig. 6. Consistent with the dominant contribution of GluN2B to the total NMDAR current, *Grin2B* appeared to be the most abundant *Grin* transcript for both controls and hibernators, along with *Grin1* that codes for the obligatory GluN1 subunit. Pharmacologically, QZN 46 is selective for both GluN2C and GluN2D. However, most neurons lacked mRNA expression of *Grin2D* (Fig 6), indicating the GluN2C is the main functional subunit blocked by QZN 46 in control neurons. Consistent with our functional results, after hibernation we observed a significant decrease in the mean abundance *Grin2C* (p=0.0003, Mann-Whitney U test). Surprisingly, although we did not observe a contribution from the GluN3 subunit to the NMDAR in either group, we observed mRNA expression of both *Grin3A* and *Grin3B*, and each of these subunits decreased after hibernation (*Grin3A*: t_34_=2.920, p=0.0062, unpaired t test; *Grin3B*: t_21.62_=2.597, p=0.0166, Welch’s t-test). These data suggest that a transcriptional program influences the NMDAR subunit composition in concert with modification of whole-cell receptor current properties.

**Figure 6.**
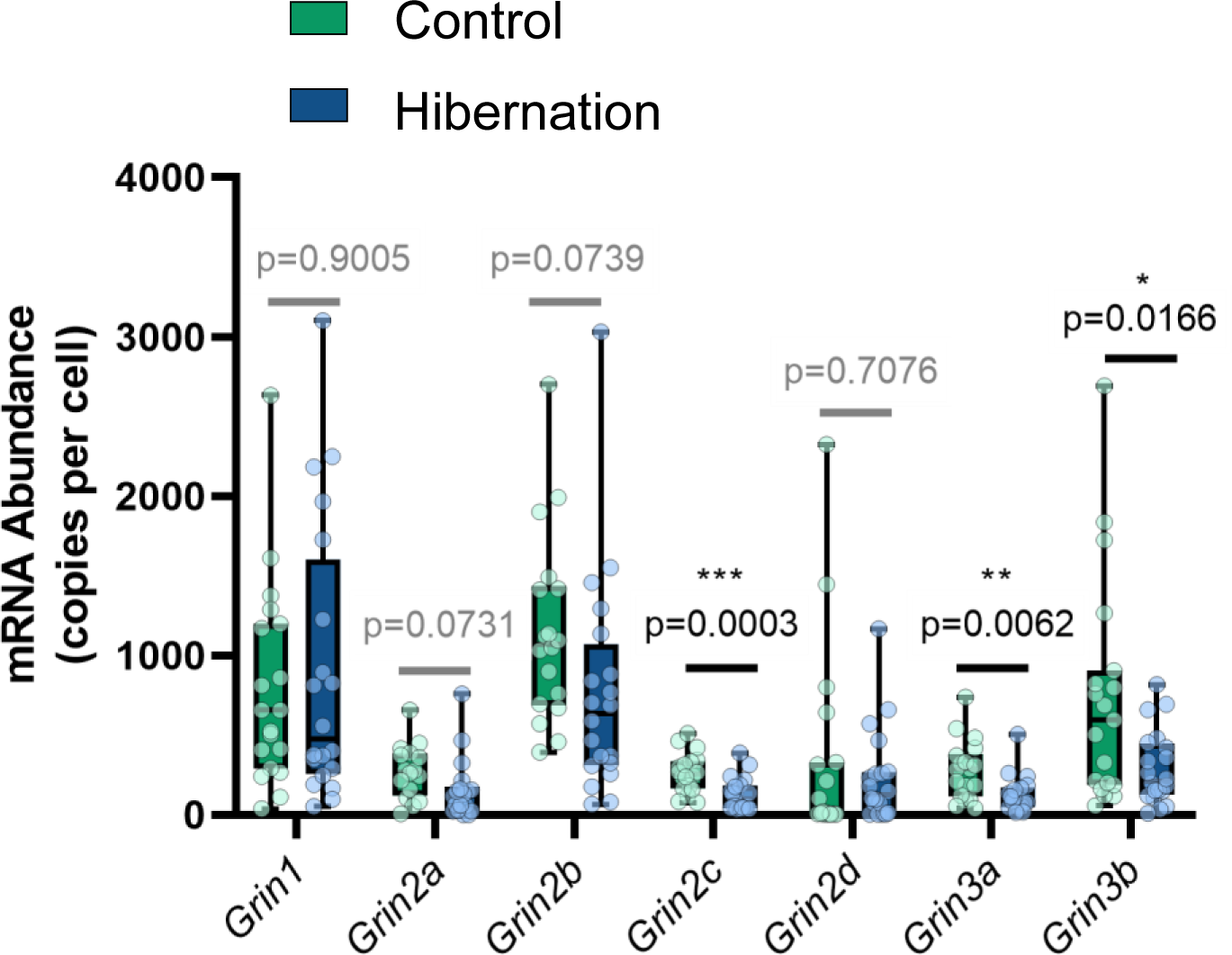
Single-cell qPCR shows that hibernation changes in the NMDAR subunit profile in motoneurons. Single-cell mRNA abundances for all 7 *Grin* family members. The dominant subunits were *Grin1*, which encodes the obligatory NMDAR subunit, GluN1, as well *Grin2B* and *Grin3B.* Hibernation led to decreases in the abundance of *Grin2C*, *Grin3A*, and *Grin3B.* Sample sizes for each gene are as follows for control and hibernation. *Grin1,* n=19 control, n=20 hibernation; *Grin2A*, n=16 control, n=16 hibernation; *Grin2B*, n=18 control, n=20 hibernation; *Grin2C,* n=19 control, n=20 hibernation; *Grin2D,* n=19 control, n=20 hibernation; *Grin3A,* n=18 control, n=18 hibernation; *Grin3B*; n=19 control, n=20 hibernation. Control cells came from N=6 animals and Hibernation cells came from N=5 animals. * p<0.05; **p<0.01; ****; p<0.001.

What are the functional impacts of NMDAR modifications on the ability of the network to function during hypoxia? In control brainstems, the first several minutes of hypoxia leads to large, uncoordinated output that interferes with the normal patterned activity of the respiratory network (Fig. 7A_1_). This type of chaotic activity is thought to arise, at least in part, due to motor hyperexcitability induced by energy stress, as it does not occur when the brainstem is well-oxygenated (Adams et al., 2021; Bueschke et al., 2021). To determine the extent to which NMDARs play a role in this response, we exposed a group of brainstem preparations to D-AP5 and then measured the degree of large amplitude output that disrupted normal activity shortly after the onset of hypoxia. In the presence of D-AP5 (Fig. 7A_2_), the mean amplitude of this chaotic “non-respiratory” activity was strongly blunted relative to controls. In addition, the small degree of disruption was often not sufficient to disrupt patterned output during hypoxia as shown in Fig. 7A_2_. These results show that the normal profile of NMDARs (with a permeability to Ca^2+^ and relatively less desensitization) contribute to chaotic output that disrupts normal function of the network during hypoxia.

**Figure 7.**
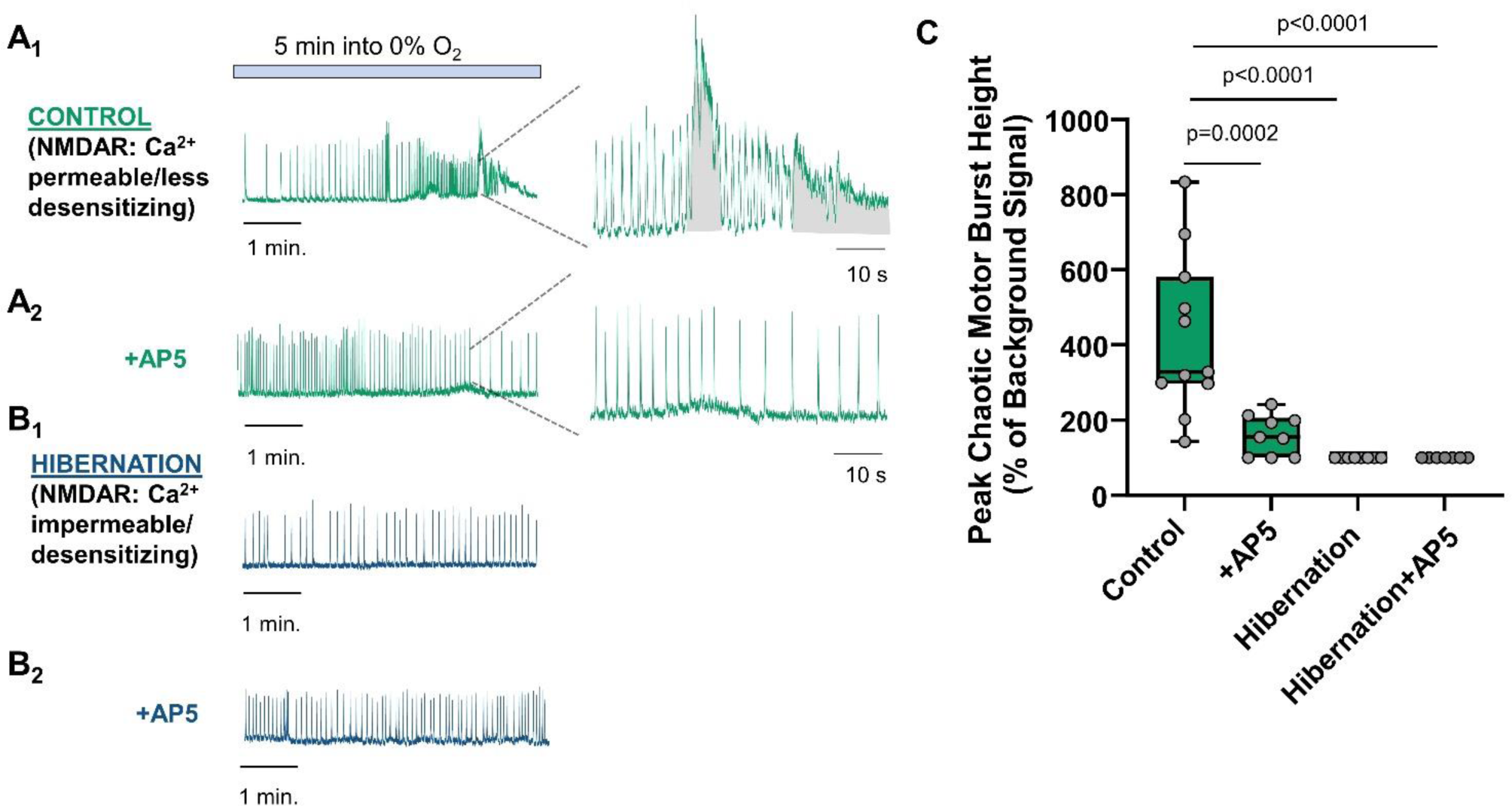
Modified NMDAR profile after hibernation protects against network hyperexcitability during hypoxia. Extracellular recordings of motor nerve output from the brainstem-spinal cord preparation. (A_1_) Shows the typical response to hypoxia in controls, whereby large amplitude bursting disrupts patterned output from the respiratory network. (A_2_) shows motor output during hypoxia in a preparation that was pretreated with D-AP5 to block NMDARs. In the presence of D-AP5, large amplitude bursting is less severe and often does not fully disrupt the respiratory rhythm. This indicates the normal profile of NMDARs contributes to disruptive motor activity in hypoxia. (B_1_) Shows patterned motor output from a hibernation preparation over the same timescale as controls. Despite intact NMDARs, there is no observable disruptive motor output. (B_2_) A hibernation preparation pretreated with D-AP5, also showing disrupted motor output. (C) Summary data of the degree of large amplitude non-respiratory bursting from n=11 controls, n=9 control+D-AP5, n=8 hibernation, and n=7 hibernation+D-AP5 analyzed by a one-way ANOVA. Non-respiratory bursting was smaller on average after the application of D-AP5, and this was not statistically different from hibernation preparations with or without D-AP5.

As disruptive motor activity during hypoxia arose largely due to NMDARs, this type of activity provided a sensitive indicator of how the modified NMDAR profile influenced network excitability in hypoxia. Strikingly, the chaotic, large amplitude, non-respiratory bursting in hypoxia that was caused largely by NMDARs did not occur in any hibernation preparation (Fig. 7B_1_). We also applied D-AP5 to a group of hibernation preparations. The addition of D-AP5 did not influence network function during hypoxia, as there was no disruptive activity to suppress (Fig. 7B_2_). These results are summarized in Fig. 7C by a one-way ANOVA for control, control+D-AP5, hibernation, and hibernation+D-AP5 (F_(3,31)_=15.25, p<0.0001, one-way ANOVA). Post hoc tests reveal that blocking NMDARs in control preparations suppresses the large non-respiratory bursting in hypoxia (p=0.0002, Holm-Sidak multiple comparisons test). This is not different than the degree of large amplitude bursting during hypoxia from hibernation preparations without (p=0.6824, Holm-Sidak multiple comparisons test) or with D-AP5 (p=0.6824, Holm-Sidak multiple comparisons test). These results show that control preparations without functional NMDARs (because they have been blocked by D-AP5) behave similar to circuits with intact NMDARs after they have become less permeable to Ca^2+^ and desensitize more strongly. Thus, modifications in NMDAR function act in a way to prevent hyperexcitable network states during energy stress while maintaining their role in network output under well-oxygenated conditions.

## Discussion

During energy stress, NMDAR activation disrupts circuit output in a wide range of species, including amphibians (Fig 7). To emerge from hibernation in an ice-covered pond, frogs must restart critical behaviors, including breathing, on the background of extraordinarily low oxygen levels (Tattersall & Ultsch, 2008; Ultsch et al., 2004). The challenge lies in the fact that most frogs are not generally considered to be a strongly “hypoxia-tolerant” species. Metabolically, a suite of plasticity mechanisms arise during hibernation and improve the fuel supply needed to operate brainstem synapses under anaerobic conditions (Bueschke et al., 2021a; Hu & Santin, 2022, Amaral-Silva & Santin, 2023). Here, we tested the hypothesis that these animals also use physiological mechanisms to avoid hyperexcitable network states induced by energy stress. We identified two key modifications to NMDARs that offset hyperexcitable output in hypoxia: reduced Ca^2+^ permeability (Fig. 3) and enhanced desensitization (Fig. 4).

These NMDAR modifications are consistent with plasticity that acts to maintain network excitability and reduce energy consumption in hypoxia. First, enhancing desensitization reduces depolarizing drive during repetitive stimulation that normally occurs during hypoxia (Fig. 4). As NMDARs contribute to chaotic bursting that disrupts patterned network output (Fig. 4A, Fig. 7A), stronger desensitization during high activity states likely acts as a potent brake on excitability. Second, switching to NMDARs with lower Ca^2+^ permeability likely incurs a cost savings. Here, roughly 50% of the population motor output that drives breathing is generated through NMDARs (Fig 1), which means for every breath the animal takes, increases in intracellular Ca^2+^ from NMDARs must be cleared to maintain ion homeostasis. Although the regulation of intracellular Ca^2+^ is multifaceted, the plasma membrane Ca^2+^ ATPase plays a large role in maintaining Ca^2+^ homeostasis in active neurons (Malci et al., 2022; Schmidt et al., 2017). This pump consumes more than 3 times the ATP as the Na^+^/K^+^ ATPase, as it extrudes 1 Ca^2+^ ion per ATP hydrolyzed and transports Ca^2+^ against a larger electrochemical gradient compared to Na^+^ and K^+^. Thus, in addition to minimizing well-established pathways for Ca^2+^-induced excitotoxicity (Szydlowska and Tymianski, 2010), lowering the Ca^2+^ permeability of NMDAR may also reduce the metabolic burden of Ca^2+^ regulation in hypoxia, allowing neurons to allocate energy to other processes required to maintain homeostasis.

Interestingly, these modifications do not alter the normal role of the NMDARs in generating motor output under well-oxygenated conditions (Fig. 1). One reason for this may be that NMDAR currents in this study were measured at the cell body. Thus, the plasticity in NMDAR function we observe may be localized to extra-synaptic regions, which are well known to elicit the pathological actions of NMDARs during hypoxia and ischemia (Tu et al., 2010; Zhou et al., 2013). It is not yet known if hibernation influences NMDARs at respiratory-related synapses. However, if hibernation alters the properties of synaptic NMDARs, reduced Ca^2+^ permeability and enhanced desensitization likely have little influence on network output since these modifications do not alter the whole-cell current amplitude and kinetics over the timescale of individual networks population bursts (<1 s burst every ∼10 s) (Fig. 1A). Altogether, NMDAR plasticity appears to contribute to a state of resilience during metabolic stress without influencing normal functioning of the network under healthy conditions.

What cellular mechanisms influence the functional changes we observe? NMDAR receptors are heterotetrameric complexes composed of an obligatory GluN1 subunit and GluN2A-D and GluN3A-B subunits, which determine receptor properties (Paoletti et al., 2013). Heterologous expression of GluN3 with GluN1 renders the receptor impermeable to Ca^2+^ and highly responsive to glycine (Chatterton et al., 2002). Although our single-cell qPCR data show altered mRNA expression of genes that code for the GluN3 subunit, our functional studies show that it does not contribute to the NMDAR current, indicating that changes in GluN3 are not responsible for decreases in Ca^2+^ permeability after hibernation. Interestingly, in NMDAR complexes containing GluN2, Ca^2+^ permeability is not an intrinsic property of the protein, but rather, is caused by phosphorylation of the GluN2B subunit (Skeberdis et al., 2006; Murphy et al., 2014). As GluN2B had the greatest functional role and mRNA expression (Fig. 5 and Fig. 6), we speculate that alterations in the phosphorylation state of GluN2B may regulate Ca^2+^ permeability. For desensitization, many factors influence this property, including subunit composition (Vicini et al., 1998), protein binding partners (Sornarajah et al., 2008), and intracellular signaling pathways (Alagarsamy et al., 1999). Most relevant to our results, GluN1/GluN2C heterodimers undergo relatively weak desensitization (Dravid et al., 2008; Alsaloum et al., 2016). We observed a decrease in *Grin2C* mRNA expression and a reduced contribution of GluN2C to the whole-cell current after hibernation. These results suggest that transcriptional control of GluN2C expression may shift neurons to favor subunit combinations with greater degrees of desensitization, such as GluN1/GluN2B (Vicini et al., 1998). As a caveat, we acknowledge that attributes of NMDAR currents may represent the sum of many potential combinations of GluN subunits; although we did not find significant functions of GluN3 and GluN2A, some individual cells within the population did appear to respond to their inhibitors, suggesting that variable combinations of NMDAR subunits compose the total whole cell current. In addition, receptor expression may vary depending on the localization within the cell (e.g., synaptic vs. extrasynaptic). This complicates the connection between single cell mRNA abundances and NMDAR function measured at the cell body. Nevertheless, combining functional and molecular approaches point to mechanisms that alter the functional properties of the NMDARs to improve function during energy stress.

The frog brainstem has a remarkable ability to improve its function during hypoxia and ischemia after hibernation (Bueschke et al., 2021). This response involves enhancing anaerobic glucose metabolism at active synapses (Amaral-Silva & Santin, 2023; Bueschke et al., 2021a). Dominant hypotheses for neural circuit evolution posit that natural selection acts on the efficiency of synapses, optimizing the ratio of information transfer to ATP consumption (Harris et al., 2015; Quintela-López et al., 2022). However, synapses here can switch into a state that supports network activity for hours with ∼1/15^th^ the amount of ATP through adjustments in synaptic metabolism that maximize glycolytic ATP production (Amaral-Silva & Santin, 2023). Our results suggest that physiological modifications help to lower the cost of network activity to match the dramatically reduced rate of energy production while maintaining seemingly normal output. Given that this circuit makes physiological and energetic modifications that allow it to run on very little energy, these results demonstrate some neural circuits may reside far from their “optimal” efficiency. This begs the question of why animals would suppress such a state until it is critical for survival (*e.g.,* emerging from an ice-covered pond after months of hibernation).

Although our results show strong support for NMDAR plasticity as a potential energy-saving mechanism, they also hint at a potential cost of maintaining an “ultra-high” efficiency network state. NMDARs play a key role in plasticity, whereby Ca^2+^ influx through the receptor potentiates synaptic strength in a wide range of systems, including the respiratory motoneurons in this species (Bueschke et al., 2021b). NMDAR modifications we observe likely incur an energy savings but have features that appear to be incompatible with NMDAR-dependent plasticity: strongly reduced Ca^2+^ permeability and greatly enhanced desensitization which quickly reduces channel activity upon repetitive stimulation. Indeed, Ca^2+^ impermeable NMDARs cause memory impairments and reduce long-term potentiation in mice (Conde-Dusman et al., 2021; Hurley et al., 2022). Therefore, we suggest a trade-off exists between metabolic robustness and the capacity for plasticity within circuits. It seems sensible for an animal to favor an extreme degree of energetic resilience to restart critical behaviors in severely hypoxic conditions. However, when animals are not faced with energy stress, as is the case for most vertebrates most of the time, it is clearly beneficial for the brain to adapt to changes in the environment through plasticity. In support of the relationship between plasticity and metabolic resilience, the rodent hippocampus shows a dorsal-ventral gradient in damage by ischemia (Ashton et al., 1989), which mirrors the capacity for synaptic plasticity owing to the degree of Ca^2+^-permeability of the NMDARs (Hurley et al., 2022). Therefore, hypotheses for selective pressures shaping the evolution of neural circuits may need to incorporate how synapses have balanced the need for plasticity and robust function during energy stress.

In conclusion, a failure to balance energy supply and physiological demands of neurons leads to disordered circuit activity. We identified a circuit that has the capacity to modify NMDARs to reduce Ca^2+^ influx and constrain excitability in hypoxia without altering their normal contribution to network function. These findings represent a state-dependent form of NMDAR plasticity that likely plays an adaptive role by promoting coordinated network activity in a circuit that cannot typically maintain homeostasis in low oxygen. These results provide insight into natural mechanisms that reduce neural circuit reliance on high rates of energy production. Yet, they also point to a potential trade-off between energetic robustness and plasticity that requires Ca^2+^ influx through NMDARs. Uncovering mechanisms that shift the balance between these two states may inform novel neuroprotection strategies with high clinical relevance.

## Author Contributions

Conception and Design; JMS. Performed Experiments; NB, MH, LAS, AA. Analyzed Data; NB, JMS. Wrote Manuscript; JMS, NB. Edited manuscript; JS, NB, MH, LAS, AA. Approved Final version of the manuscript; JS, NB, MH, LAS, AA.

## Acknowledgment

We would like to thank the National Institutes of Health (R15NS112920-01A1, R01NS114514) the U.S. Department of Defense (W911NF2010275) for funding to JMS.

## References

Abrahamsson T, Chou CYC, Li SY, Mancino A, Costa RP, Brock JA, Nuro E, Buchanan KA, Elgar D, Blackman AV (2017). Differential regulation of evoked and spontaneous release by presynaptic NMDA receptors. Neuron 96:839–855. e835. 10.1016/j.neuron.2017.09.030

Adams, S., Zubov, T., Bueschke, N., & Santin, J. M. (2021). Neuromodulation or energy failure? Metabolic limitations silence network output in the hypoxic amphibian brainstem. American Journal of Physiology-Regulatory, Integrative and Comparative Physiology, 320(2), R105–R116. 10.1152/ajpregu.00209.2020

Alagarsamy, S., Marino, M. J., Rouse, S. T., Gereau, R. W., Heinemann, S. F., & Conn, P. J. (1999). Activation of NMDA receptors reverses desensitization of mGluR5 in native and recombinant systems. Nature Neuroscience, 2(3), 234–240. 10.1038/6338

Alsaloum M, Kazi R, Gan Q, Amin J, Wollmuth LP (2016). A molecular determinant of subtype-specific desensitization in ionotropic glutamate receptors. Journal of Neuroscience 36, 2617–2622. 10.1523/JNEUROSCI.2667-15.2016

Amaral-Silva, L., Santin, J.M., (2023). Synaptic modifications transform neural networks to function without oxygen. BMC Biology, 21(54).

Ashton, D., Van Reempts, J., Haseldonckx, M., & Willems, R. (1989). Dorsal-ventral gradient in vulnerability of CA1 hippocampus to ischemia: A combined histological and electrophysiological study. Brain Research, 487(2), 368–372. 10.1016/0006-8993(89)90842-1

Beesley, S., Sullenberger, T., & Kumar, S. S. (2020). The GluN3 subunit regulates ion selectivity within native N-methyl-d-aspartate receptors. IBRO Reports, 9, 147–156. 10.1016/j.ibror.2020.07.009

Bettini E, Sava A, Griffante C, Carignani C, Buson A, Capelli AM, Negri M, Andreetta F, Senar-Sancho SA, Guiral L (2010). Identification and characterization of novel NMDA receptor antagonists selective for NR2A-over NR2B-containing receptors. Journal of Pharmacology and Experimental Therapeutics 335, 636–644. 10.1124/jpet.110.172544

Bickler, P. E., & Buck, L. T. (2007). Hypoxia Tolerance in Reptiles, Amphibians, and Fishes: Life with Variable Oxygen Availability. Annual Review of Physiology, 69(1), 145–170. 10.1146/annurev.physiol.69.031905.162529

Bickler, P. E., Donohoe, P. H., & Buck, L. T. (2000). Hypoxia-Induced Silencing of NMDA Receptors in Turtle Neurons. The Journal of Neuroscience, 20(10), 3522– 3528. 10.1523/JNEUROSCI.20-10-03522.2000

Bordone, M. P., Salman, M. M., Titus, H. E., Amini, E., Andersen, J. V., Chakraborti, B., Diuba, A. V., Dubouskaya, T. G., Ehrke, E., Espindola de Freitas, A., Braga de Freitas, G., Gonçalves, R. A., Gupta, D., Gupta, R., Ha, S. R., Hemming, I. A., Jaggar, M., Jakobsen, E., Kumari, P., … Seidenbecher, C. I. (2019). The energetic brain – A review from students to students. Journal of Neurochemistry, 151(2), 139–165. 10.1111/jnc.14829

Buck, L. T., & Pamenter, M. E. (2018). The hypoxia-tolerant vertebrate brain: Arresting synaptic activity. Comparative Biochemistry and Physiology Part B: Biochemistry and Molecular Biology, 224, 61–70. 10.1016/j.cbpb.2017.11.015

Bueschke, N., Amaral-Silva, L., Adams, S., & Santin, J. M. (2021a). Transforming a neural circuit to function without oxygen and glucose delivery. Current Biology, 31(24), R1564–R1565. 10.1016/j.cub.2021.11.003

Bueschke, N., Amaral-Silva, L., Hu, M., & Santin, J. M. (2021b). Lactate ions induce synaptic plasticity to enhance output from the central respiratory network. The Journal of Physiology, 599(24), 5485–5504. 10.1113/JP282062

Chatterton, J. E., Awobuluyi, M., Premkumar, L. S., Takahashi, H., Talantova, M., Shin, Y., Cui, J., Tu, S., Sevarino, K. A., Nakanishi, N., Tong, G., Lipton, S. A., & Zhang, D. (2002). Excitatory glycine receptors containing the NR3 family of NMDA receptor subunits. Nature, 415(6873), 793–798. 10.1038/nature715

Christian DT, Stefanik MT, Bean LA, Loweth JA, Wunsch AM, Funke JR, Briggs CA, Lyons J, Neal D, Milovanovic M (2021). GluN3-containing NMDA receptors in the rat nucleus accumbens core contribute to incubation of cocaine craving. Journal of Neuroscience 41, 8262–8277. 10.1523/JNEUROSCI.0406-21.2021

Conde-Dusman, M. J., Dey, P. N., Elía-Zudaire, Ó., Rabaneda, L. G., García-Lira, C., Grand, T., Briz, V., Velasco, E. R., Andero, R., Niñerola, S., Barco, A., Paoletti, P., Wesseling, J. F., Gardoni, F., Tavalin, S. J., & Perez-Otaño, I. (2021). Control of protein synthesis and memory by GluN3A-NMDA receptors through inhibition of GIT1/mTORC1 assembly. ELife, 10, e71575. 10.7554/eLife.71575

Czech-Damal, N. U., Geiseler, S. J., Hoff, M. L. M., Schliep, R., Ramirez, J.-M., Folkow, L. P., & Burmester, T. (2014). The role of glycogen, glucose and lactate in neuronal activity during hypoxia in the hooded seal (Cystophora cristata) brain. Neuroscience, 275, 374–383. 10.1016/j.neuroscience.2014.06.024

Dravid, S. M., Prakash, A., & Traynelis, S. F. (2008). Activation of recombinant NR1/NR2C NMDA receptors: NR1/NR2C receptor activation. The Journal of Physiology, 586(18), 4425–4439. 10.1113/jphysiol.2008.158634

Edman S, McKay S, Macdonald L, Samadi M, Livesey M, Hardingham G, Wyllie D (2012). TCN 201 selectively blocks GluN2A-containing NMDARs in a GluN1 co-agonist dependent but non-competitive manner. Neuropharmacology 63, 441–449. 10.1016/j.neuropharm.2012.04.027

France G, Fernández-Fernández D, Burnell ES, Irvine MW, Monaghan DT, Jane DE, Bortolotto ZA, Collingridge GL, Volianskis A (2017). Multiple roles of GluN2B-containing NMDA receptors in synaptic plasticity in juvenile hippocampus. Neuropharmacology 1127 6–83. 10.1016/j.neuropharm.2016.08.010

Garcia, V. B., Abbinanti, M. D., Harris-Warrick, R. M., & Schulz, D. J. (2018). Effects of Chronic Spinal Cord Injury on Relationships among Ion Channel and Receptor mRNAs in Mouse Lumbar Spinal Cord. Neuroscience, 393, 42–60. 10.1016/j.neuroscience.2018.09.034

Hammond, S. A., Warren, R. L., Vandervalk, B. P., Kucuk, E., Khan, H., Gibb, E. A., Pandoh, P., Kirk, H., Zhao, Y., Jones, M., Mungall, A. J., Coope, R., Pleasance, S., Moore, R. A., Holt, R. A., Round, J. M., Ohora, S., Walle, B. V., Veldhoen, N.,… Birol, I. (2017). The North American bullfrog draft genome provides insight into hormonal regulation of long noncoding RNA. Nature Communications, 8(1), 1433. 10.1038/s41467-017-01316-7

Hansen KB, Traynelis SF (2011). Structural and mechanistic determinants of a novel site for noncompetitive inhibition of GluN2D-containing NMDA receptors. Journal of Neuroscience 31, 3650–3661. 10.1523/JNEUROSCI.5565-10.2011

Harris, J. J., Jolivet, R., Engl, E., & Attwell, D. (2015). Energy-Efficient Information Transfer by Visual Pathway Synapses. Current Biology, 25(24), 3151–3160. 10.1016/j.cub.2015.10.063

Hille, B. (2001). *Ion channels of excitable membranes* (3rd ed). Sinauer.

Howell, B., Baumgardner, F., Bondi, K., & Rahn, H. (1970). Acid-base balance in cold-blooded vertebrates as a function of body temperature. American Journal of Physiology-Legacy Content, 218(2), 600–606. 10.1152/ajplegacy.1970.218.2.600

Hu, M., & Santin, J. M. (2022). Transformation to ischaemia tolerance of frog brain function corresponds to dynamic changes in mRNA co-expression across metabolic pathways. Proceedings of the Royal Society B: Biological Sciences, 289(1979), 20221131. 10.1098/rspb.2022.1131

Hurley, E. P., Mukherjee, B., Fang, L., Barnes, J. R., Nafar, F., Hirasawa, M., & Parsons, M. P. (2022). *Atypical NMDA Receptors Limit Synaptic Plasticity in the Adult Ventral Hippocampus* [Preprint]. Neuroscience. 10.1101/2022.10.05.510966

Jatzke, C., Junryo Watanabe, & Wollmuth, L. P. (2002). Voltage and concentration dependence of Ca ^2+^ permeability in recombinant glutamate receptor subtypes. The Journal of Physiology, 538(1), 25–39. 10.1113/jphysiol.2001.012897

Kottick, A., Baghdadwala, M. I., Ferguson, E. V., & Wilson, R. J. A. (2013). Transmission of the respiratory rhythm to trigeminal and hypoglossal motor neurons in the American Bullfrog (Lithobates catesbeiana). Respiratory Physiology & Neurobiology, 188(2), 180–191. 10.1016/j.resp.2013.06.008

Kvist T, Greenwood JR, Hansen KB, Traynelis SF, Bräuner-Osborne H (2013). Structure-based discovery of antagonists for GluN3-containing N-methyl-D-aspartate receptors. Neuropharmacology 75:324–336. 10.1016/j.neuropharm.2013.08.003

Larson, J., Drew, K. L., Folkow, L. P., Milton, S. L., & Park, T. J. (2014). No oxygen? No problem! Intrinsic brain tolerance to hypoxia in vertebrates. Journal of Experimental Biology, 217(7), 1024–1039. 10.1242/jeb.085381

Larson, J., & Park, T. J. (2009). Extreme hypoxia tolerance of naked mole-rat brain. NeuroReport, 20(18), 1634–1637. 10.1097/WNR.0b013e32833370cf

Malci, A., Lin, X., Sandoval, R., Gundelfinger, E. D., Naumann, M., Seidenbecher, C. I., & Herrera-Molina, R. (2022). Ca2+ signaling in postsynaptic neurons: Neuroplastin-65 regulates the interplay between plasma membrane Ca2+ ATPases and ionotropic glutamate receptors. Cell Calcium, 106, 102623. 10.1016/j.ceca.2022.102623

Murphy, J. A., Stein, I. S., Lau, C. G., Peixoto, R. T., Aman, T. K., Kaneko, N., Aromolaran, K., Saulnier, J. L., Popescu, G. K., Sabatini, B. L., Hell, J. W., & Zukin, R. S. (2014). Phosphorylation of Ser1166 on GluN2B by PKA Is Critical to Synaptic NMDA Receptor Function and Ca ^2+^ Signaling in Spines. The Journal of Neuroscience, 34(3), 869–879. 10.1523/JNEUROSCI.4538-13.2014

Paoletti, P., Bellone, C., & Zhou, Q. (2013). NMDA receptor subunit diversity: Impact on receptor properties, synaptic plasticity and disease. Nature Reviews Neuroscience, 14(6), 383–400. 10.1038/nrn3504

Pellizzari, S., Hu, M., Amaral-Silva, L., Saunders, S., Santin, J.M. (2023). Neuron populations use variable combinations of short-term feedback mechanisms to stabilize firing rate. PLOS Biology, accepted for publication.

Quintela-López, T., Shiina, H., & Attwell, D. (2022). Neuronal energy use and brain evolution. Current Biology, 32(12), R650–R655. 10.1016/j.cub.2022.02.005

Santin, J. M., & Schulz, D. J. (2019). Membrane Voltage Is a Direct Feedback Signal That Influences Correlated Ion Channel Expression in Neurons. Current Biology, 29(10), 1683–1688.e2. 10.1016/j.cub.2019.04.008

Schmidt, N., Kollewe, A., Constantin, C. E., Henrich, S., Ritzau-Jost, A., Bildl, W., Saalbach, A., Hallermann, S., Kulik, A., Fakler, B., & Schulte, U. (2017). Neuroplastin and Basigin Are Essential Auxiliary Subunits of Plasma Membrane Ca2+-ATPases and Key Regulators of Ca2+ Clearance. Neuron, 96(4), 827–838.e9. 10.1016/j.neuron.2017.09.038

Skeberdis, V. A., Chevaleyre, V., Lau, C. G., Goldberg, J. H., Pettit, D. L., Suadicani, S. O., Lin, Y., Bennett, M. V. L., Yuste, R., Castillo, P. E., & Zukin, R. S. (2006). Protein kinase A regulates calcium permeability of NMDA receptors. Nature Neuroscience, 9(4), 501–510. 10.1038/nn1664

Sornarajah, L., Vasuta, O. C., Zhang, L., Sutton, C., Li, B., El-Husseini, A., & Raymond, L. A. (2008). NMDA Receptor Desensitization Regulated by Direct Binding to PDZ1-2 Domains of PSD-95. Journal of Neurophysiology, 99(6), 3052–3062. 10.1152/jn.90301.2008

Szydlowska, K., & Tymianski, M. (2010). Calcium, ischemia and excitotoxicity. Cell Calcium, 47(2), 122–129. 10.1016/j.ceca.2010.01.003

Tattersall, G. J., & Boutilier, R. G. (1997). Balancing hypoxia and hypothermia in cold-submerged frogs. The Journal of Experimental Biology, 200(Pt 6), 1031–1038. 10.1242/jeb.200.6.1031

Tattersall, G. J., & Ultsch, G. R. (2008). Physiological Ecology of Aquatic Overwintering in Ranid Frogs. Biological Reviews, 83(2), 119–140. 10.1111/j.1469-185X.2008.00035.x

Tu, W., Xu, X., Peng, L., Zhong, X., Zhang, W., Soundarapandian, M. M., Belal, C., Wang, M., Jia, N., Zhang, W., Lew, F., Chan, S. L., Chen, Y., & Lu, Y. (2010). DAPK1 Interaction with NMDA Receptor NR2B Subunits Mediates Brain Damage in Stroke. Cell, 140(2), 222–234. 10.1016/j.cell.2009.12.055

Ultsch, G. R., Reese, S. A., & Stewart, E. R. (2004). Physiology of hibernation inRana pipiens: Metabolic rate, critical oxygen tension, and the effects of hypoxia on several plasma variables. Journal of Experimental Zoology, 301A(2), 169–176. 10.1002/jez.a.20014

Vicini, S., Wang, J. F., Li, J. H., Zhu, W. J., Wang, Y. H., Luo, J. H., Wolfe, B. B., & Grayson, D. R. (1998). Functional and Pharmacological Differences Between Recombinant *N*-Methyl-D-Aspartate Receptors. Journal of Neurophysiology, 79(2), 555–566. 10.1152/jn.1998.79.2.555

White, R. J., & Reynolds, I. J. (1995). Mitochondria and Na+/Ca2+ exchange buffer glutamate-induced calcium loads in cultured cortical neurons. The Journal of Neuroscience: The Official Journal of the Society for Neuroscience, 15(2), 1318– 1328.

Wilkie, M. P., Pamenter, M. E., Alkabie, S., Carapic, D., Shin, D. S. H., & Buck, L. T. (2008). Evidence of anoxia-induced channel arrest in the brain of the goldfish (Carassius auratus). Comparative Biochemistry and Physiology Part C: Toxicology & Pharmacology, 148(4), 355–362. 10.1016/j.cbpc.2008.06.004

Wu, Q. J., & Tymianski, M. (2018). Targeting NMDA receptors in stroke: New hope in neuroprotection. Molecular Brain, 11(1), 15. 10.1186/s13041-018-0357-8

Zhou, X., Hollern, D., Liao, J., Andrechek, E., & Wang, H. (2013). NMDA receptor-mediated excitotoxicity depends on the coactivation of synaptic and extrasynaptic receptors. Cell Death & Disease, 4(3), e560–e560. 10.1038/cddis.2013.82

